# Taking population stratification into account by local permutations in rare-variant association studies on small samples

**DOI:** 10.1101/2020.01.29.924977

**Authors:** J. Mullaert, M. Bouaziz, Y. Seeleuthner, B. Bigio, J-L. Casanova, A. Alcais, L. Abel, A. Cobat

## Abstract

Many methods for rare variant association studies require permutations to assess the significance of tests. Standard permutations assume that all individuals are exchangeable and do not take population stratification (PS), a known confounding factor in genetic studies, into account. We propose a novel strategy, *LocPerm*, in which individuals are permuted only with their closest ancestry-based neighbors. We performed a simulation study, focusing on small samples, to evaluate and compare *LocPerm* with standard permutations and classical adjustment on first principal components. Under the null hypothesis, *LocPerm* was the only method providing an acceptable type I error, regardless of sample size and level of stratification. The power of *LocPerm* was similar to that of standard permutation in the absence of PS, and remained stable in different PS scenarios. We conclude that *LocPerm* is a method of choice for taking PS and/or small sample size into account in rare variant association studies.

## Introduction

Population stratification (PS) is a classic confounding factor in genetic association studies of common variants (1). It also affects association studies involving rare variants in the context of next-generation sequencing (NGS) analyses (2–4). Principal component analysis (PCA) is the most widely used approach to correction for stratification. The use of principal components (PCs) computed from common variants as covariates in a regression framework to test for association has been widely investigated (1,5,6). This strategy provides a satisfactory correction in a number of settings, but is subject to several limitations, particularly in cases of complex population structure (4,7). In addition, the regression framework implicitly assumes an asymptotic distribution of the test statistics, which is rarely achieved when sample size is small (8), and few studies of PC-based correction in this context have been published (9).

Permutation methods (particularly the derivation of an empiric distribution by the random permutation of phenotype labels) are classically used in strategies for deriving *p*-values from a test statistic with a probability distribution that is unknown or from which it is difficult to sample (10). However, this approach assumes that all individuals are equally interchangeable under the null hypothesis, an assumption that is not valid in the presence of PS (11). When ancestry is known, it is reasonable to ensure that permutations result exclusively in the exchange of individuals of the same ancestry, but this information is rarely available in practice. We investigated the impact of PS on association studies based on NGS data in the context of limited sample sizes, a situation frequently observed in rare disorders. We propose a new method, *LocPerm*, based on population-adapted permutation and taking into account the genetic distance between individuals. We describe a detailed analysis of its properties with respect to PC adjustment and standard permutation.

## Materials and methods

We propose a new approach, *LocPerm*, in which permutation is restricted such that each individual can be exchanged only with one of its nearest neighbors in terms of a PC-based genetic distance (Supplementary note). Here, we focused on a binary phenotype and the cohort allelic sum test (CAST) approach (12) implemented in a logistic regression framework, using the likelihood ratio test (LRT) statistic. The *LocPerm p*-value can be calculated by either the usual *full empiric* (*FE*) approach (in which the *p*-value is equal to the number of permutation samples with a test statistic as extreme as that observed, divided by the total number of permutation samples), or a *semi-empiric* (*SE*) approach. In the SE approach, a limited number of permuted statistics are used to estimate the mean (m) and standard deviation (*σ*) of the test statistic under the null hypothesis, and the *p*-value is calculated from the N(m, σ^2^) distribution (Supplementary note).

We performed a simulation study based on real NGS data, to assess the type I error and power of the *LocPerm* procedure in the context of small sample sizes. We compared *LocPerm* to the asymptotic CAST approach with (CAST-3PC) and without (CAST) inclusion of the three principal components (PCs) in the regression model, and to standard permutations applied to CAST. We extracted 1,523 individuals — 745 of Southern European ancestry, 651 of Central European ancestry and 127 of Northern European ancestry (**eFigure1**) — from our in-house HGID (Human Genetic of Infectious Diseases) whole-exome sequencing dataset and the public 1000 Genomes Phase 3 whole-genome sequencing dataset (Supplementary note). Under the null hypothesis, cases and controls were randomly drawn from the source population according to three PS scenarios (absence of PS, intermediate and extreme stratification, supplementary note). For power analysis, we selected one gene with a cumulative frequency of rare variants of 6.2%, for which we simulated a binary phenotype in the source population, assuming a relative risk of the disease of 4 for individuals carrying at least one rare variant. We then conducted a sensitivity analysis to investigate the effect on the type I error of the number of neighbors in the *LocPerm* procedure.

**Figure 1:**
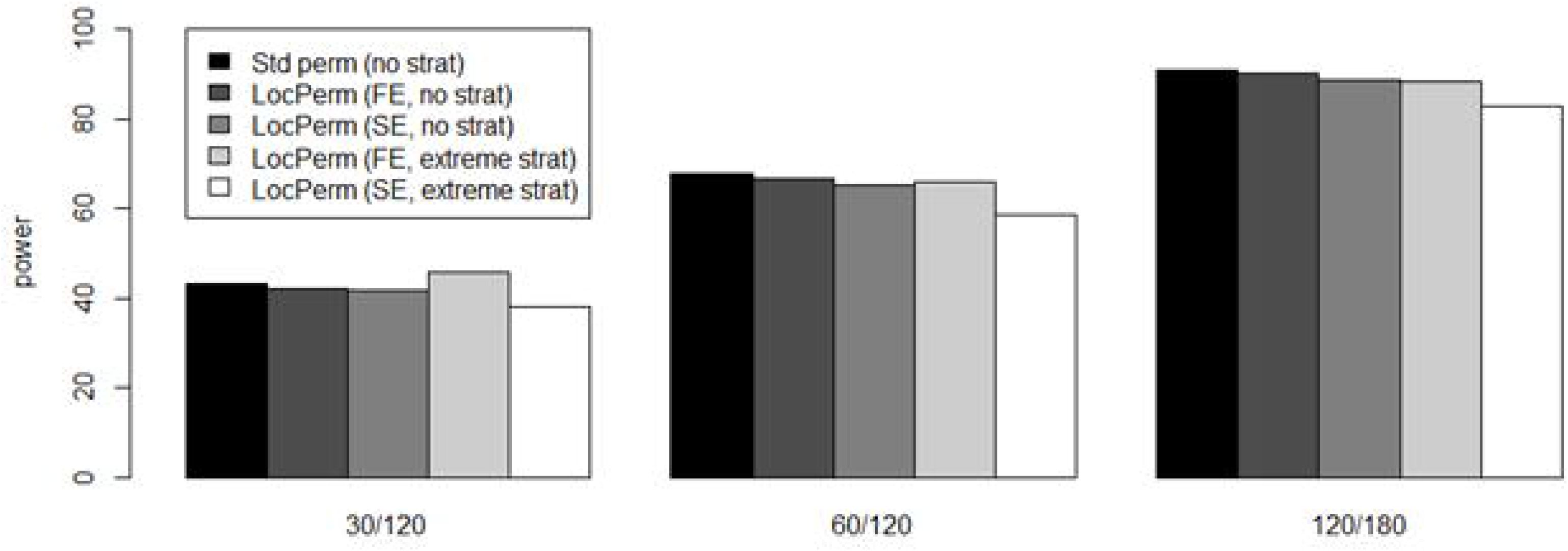
Power at 1% significance level for different PS scenarios, permutation procedures and number of cases and controls in the sample

## Results

The results of the simulation study under the null hypothesis (H_0_) for the three stratification scenarios and various sample sizes are shown in **Table 1**(for *α*=0.01) and **eTable 1**(for *α*=0.05). In the absence of PS, methods based on test statistics following an asymptotic distribution (CAST and CAST-3PC) had inflated type I errors for small sample sizes. Stronger inflation was observed for CAST-3PC than for CAST (e.g. type I error=0.0124 vs. 0.0114 at *α*=0.01 for samples of 30 cases and 180 controls). Methods based on permutation (standard and *LocPerm*) gave correct type I errors. In the presence of PS, the strongest type I error inflation was observed for CAST. The addition of the first three PCs to the model took PS into account only partially. Inflated type I errors were also observed for standard permutations in the presence of PS. Type I error inflation increased with the degree of PS and with sample size for CAST and standard permutation, whereas small sample size appeared to be the main source of inflation for CAST-3PC. The *LocPerm* procedures (*FE* and *SE*) provided type I errors close to the expected *α* threshold across all sample sizes and PS scenarios, despite being slightly conservative in the presence of extreme stratification, particularly for *LocPerm*-*SE*.

**Table 1.**
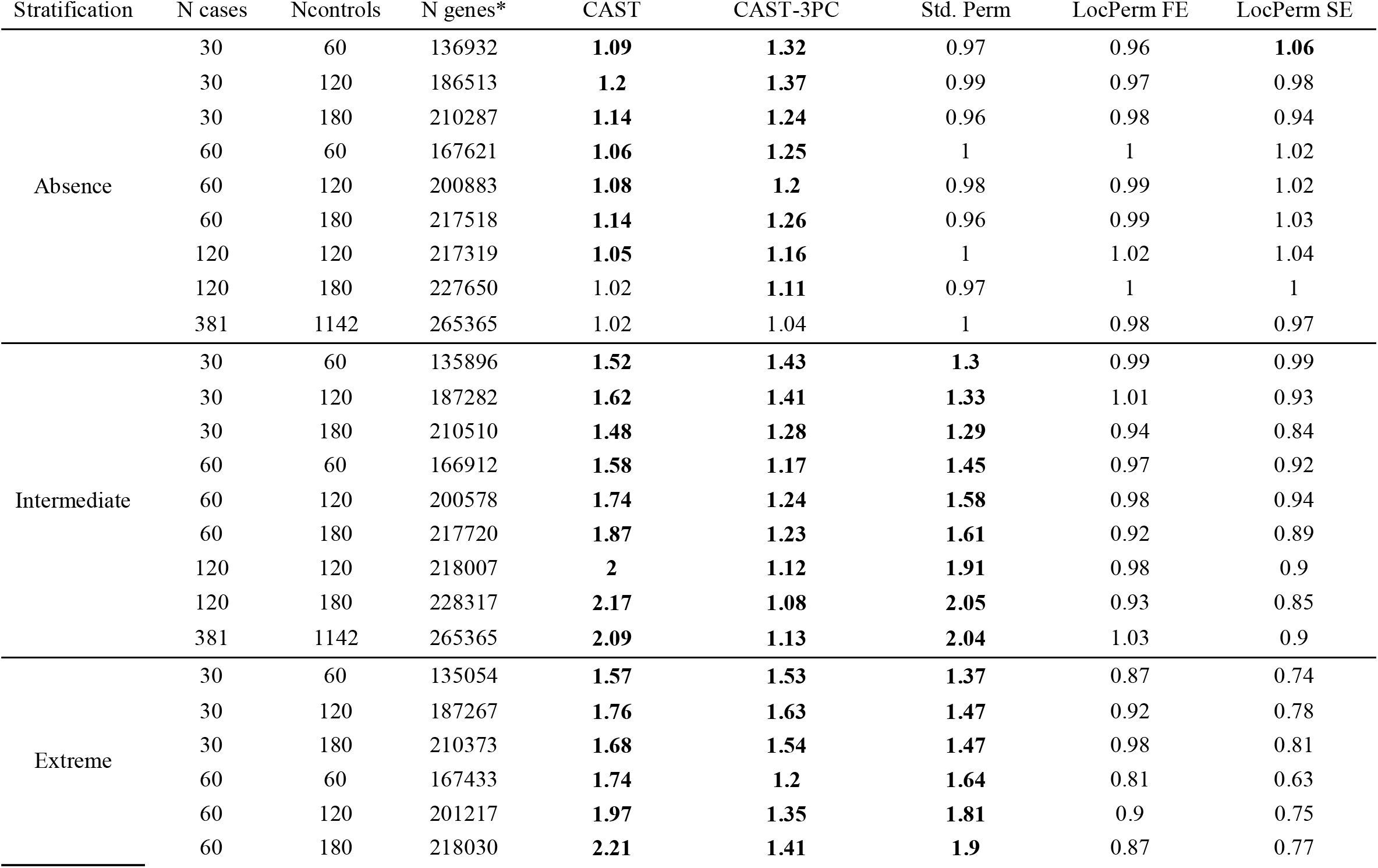
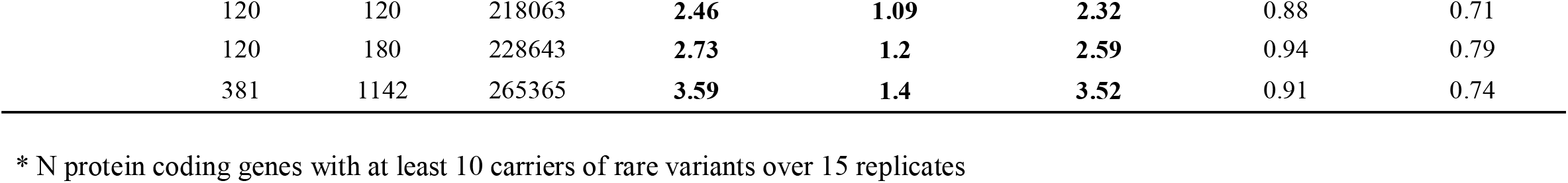
Type I error rates of the different approaches and stratification scenarios at a nominal alpha level of 1%. Type I error rates above the upper bound of the 95% prediction interval in bold.

| Stratification | N cases | Ncontrols | N genes* | CAST | CAST-3PC | Std. Perm | LocPerm FE | LocPerm SE |
| --- | --- | --- | --- | --- | --- | --- | --- | --- |
| Absence | 30 | 60 | 136932 | **1.09** | **1.32** | 0.97 | 0.96 | **1.06** |
|  | 30 | 120 | 186513 | **1.2** | **1.37** | 0.99 | 0.97 | 0.98 |
|  | 30 | 180 | 210287 | **1.14** | **1.24** | 0.96 | 0.98 | 0.94 |
|  | 60 | 60 | 167621 | **1.06** | **1.25** | 1 | 1 | 1.02 |
|  | 60 | 120 | 200883 | **1.08** | **1.2** | 0.98 | 0.99 | 1.02 |
|  | 60 | 180 | 217518 | **1.14** | **1.26** | 0.96 | 0.99 | 1.03 |
|  | 120 | 120 | 217319 | **1.05** | **1.16** | 1 | 1.02 | 1.04 |
|  | 120 | 180 | 227650 | 1.02 | **1.11** | 0.97 | 1 | 1 |
|  | 381 | 1142 | 265365 | 1.02 | 1.04 | 1 | 0.98 | 0.97 |
| Intermediate | 30 | 60 | 135896 | **1.52** | **1.43** | **1.3** | 0.99 | 0.99 |
|  | 30 | 120 | 187282 | **1.62** | **1.41** | **1.33** | 1.01 | 0.93 |
|  | 30 | 180 | 210510 | **1.48** | **1.28** | **1.29** | 0.94 | 0.84 |
|  | 60 | 60 | 166912 | **1.58** | **1.17** | **1.45** | 0.97 | 0.92 |
|  | 60 | 120 | 200578 | **1.74** | **1.24** | **1.58** | 0.98 | 0.94 |
|  | 60 | 180 | 217720 | **1.87** | **1.23** | **1.61** | 0.92 | 0.89 |
|  | 120 | 120 | 218007 | **2** | **1.12** | **1.91** | 0.98 | 0.9 |
|  | 120 | 180 | 228317 | **2.17** | **1.08** | **2.05** | 0.93 | 0.85 |
|  | 381 | 1142 | 265365 | **2.09** | **1.13** | **2.04** | 1.03 | 0.9 |
| Extreme | 30 | 60 | 135054 | **1.57** | **1.53** | **1.37** | 0.87 | 0.74 |
|  | 30 | 120 | 187267 | **1.76** | **1.63** | **1.47** | 0.92 | 0.78 |
|  | 30 | 180 | 210373 | **1.68** | **1.54** | **1.47** | 0.98 | 0.81 |
|  | 60 | 60 | 167433 | **1.74** | **1.2** | **1.64** | 0.81 | 0.63 |
|  | 60 | 120 | 201217 | **1.97** | **1.35** | **1.81** | 0.9 | 0.75 |
|  | 60 | 180 | 218030 | **2.21** | **1.41** | **1.9** | 0.87 | 0.77 |

|  |  |  |  |  |  |  |  |
| --- | --- | --- | --- | --- | --- | --- | --- |
| 120 | 120 | 218063 | **2.46** | **1.09** | **2.32** | 0.88 | 0.71 |
| 120 | 180 | 228643 | **2.73** | **1.2** | **2.59** | 0.94 | 0.79 |
| 381 | 1142 | 265365 | **3.59** | **1.4** | **3.52** | 0.91 | 0.74 |

\* N protein coding genes with at least 10 carriers of rare variants over 15 replicates

We further investigated the sensitivity of the *LocPerm* procedure to the number of neighbors under H_0_ (Figure 2). With an *α* threshold of 0.01, the type I error of the *LocPerm* procedure remained stable over a wide range of numbers of neighbors (from 20 to 170 for a total sample of 210 individuals), and the use of 30 neighbors appeared to be a reasonable choice. The results of the simulation study under the alternative hypothesis are shown in **Figure 1** for methods providing a non-inflated type I error rate (i.e. standard permutation in the absence of stratification and *LocPerm*-*SE* and *LocPerm*-*FE* with and without stratification). In the absence of stratification, a similar power was achieved for standard permutation, *LocPerm*-*FE* and *LocPerm*-*SE* (43%, 42% and 42% at *α*=0.01 for standard permutation, *LocPerm*-*FE* and *LocPerm*-*SE*, respectively). In the presence of extreme stratification, the power of *LocPerm*-*FE* was well conserved, whereas that of *LocPerm*-*SE* decreased slightly, consistent with its conservative type I error rate in this scenario.

**Figure 2:**
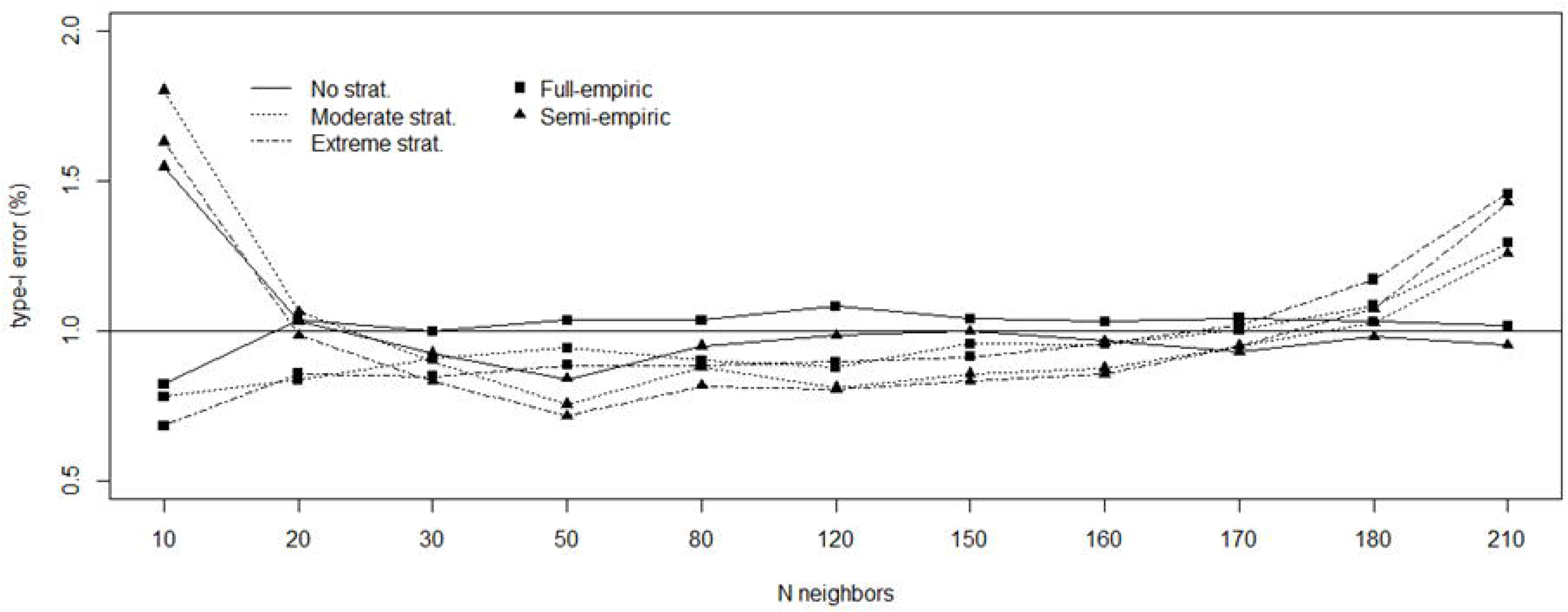
Influence of the number of neighbors for the generation of local permutation (x axis) on type I error (y axis) for the scenario with 30 cases and 180 controls. The situation with 210 neighbors corresponds to standard permutation.

## Discussion

The inclusion of the first few PCs in the association model is a popular strategy for taking population structure into account. However, it is suitable only for methods implemented in a regression framework and requires large sample sizes. We found that, in small samples, inclusion of the first three PCs in CAST failed to control the type I error in the presence of PS. In situations in which permutations were required, the *LocPerm* procedure proposed here took PS into account effectively, with no significant power loss relative to other methods in the absence of PS. The SE approximation performed well in all scenarios, being only slightly conservative in the context of extreme PS and reducing the computational cost by a factor 10 relative to the *FE* approach. We did not include adaptive permutation (13), in which the number of permutation samples decreases as the observed *p*-value increases, in our comparison. However, we would expect the *SE* approximation to be faster than adaptive permutation because it requires only 500 permutation samples, whatever the observed *p*-value.

A permutation approach handling PS was proposed in a previous study (14). The odds of disease conditional on covariates were estimated under a null model of no genetic association, and individual phenotypes were resampled, using these disease probabilities as individual weights, to obtain permuted data with a similar PS. However, subsequent studies showed that this procedure was less efficient than regular PC correction for dealing with fine-scale population structure (15). We show here that *LocPerm*, which uses the first 10 PCs weighted by their eigenvalues to compute a genetic distance matrix, handles complex and extreme PS more effectively than the standard PC-based correction approach, particularly in the context of small sample size. We focused here on binary traits and the CAST approach, but it should be straightforward to extend the *LocPerm* approach to quantitative traits and other rare variant association tests, particularly for adaptive burden tests requiring permutations.

## Acknowledgment

We thank both branches of the Laboratory of Human Genetics of Infectious Diseases for helpful discussions and support.

