## Supplementary material for "Taking population stratification into account by local permutations in rare-variant association studies on small samples"

A. Cobat

### 1. Supplementary note

#### *a. The LocPerm procedure*

Standard permutation procedure consists in computing several test statistics ($T_1$, ..., $T_B$) after phenotype permutation in order to generate an empiric distribution as close as possible to the unknown true null distribution (1). The permuted statistics are used to derive empirical p-values. However, it makes the hypothesis that all individuals are exchangeable which is not the case in the presence of population structure. Under the null hypothesis, individuals with the same ancestry are more likely to share the same phenotype than individual from different ancestries. Therefore, our idea is to establish for each sample a neighborhood, i.e. a set of relatively close samples with whom it is reasonable to exchange phenotypes, based on the genetic distance derived from the sample coordinates along principal component axes calculated on common variants. The genetic distance between two individuals i and j was computed as $d_{ij}^2 = \sum_{k=1}^{10} \lambda_k \left|PC_{ki} - PC_{kj}\right|^2$, where PC is the matrix of principal components (PCs) calculated on common variants and $\lambda_k$ the eigenvalue corresponding to the k-th principal component $PC_k$. We ended the summation at the $10^{th}$ component since the proportion of explained variance was very high and the resulting distance would be only slightly modified with additional components.

We set a number *N* and only allow permutation that ensure that each phenotype is drawn from the *N* nearest neighbors, in the sense of the genetic distance. A permutation satisfying this constraint is called a restricted (or local) permutation. To generate a list of such restricted permutation, we generated a sequence of permutation starting from the identity as a random walk in the set of restricted permutations in the following way: individual $i$ is randomly exchanged with another one (possibly himself), to ensure that the resulting permutation satisfies the original constraint. This elementary step is repeated for every individual $i$ and the resulting permutation is the next step of the random walk. This is nothing but a Gibbs sampler whose limit distribution is the uniform distribution on restricted permutations. We used a burn-in of 100 iterations (i.e. the first 100 permutations are dropped) and a step of 10 to ensure relative independence between to permutation in the list (i.e. only 1 iteration out of 10 outputs of the Gibbs sampler are considered). As shown in the results section, the procedure was rather stable within a large range of N values and we used *N=30* in the simulation study.

*b. Full- and semi-empiric p-value derivation*

P-values are usually defined as the proportion of permuted test statistics at least as extreme as the observed one. To account for a possible discrete distribution of test statistics in the context of small samples, we adapted this procedure and draw the p-value from a uniform distribution U([a,b]), where a (resp. b) stands for the observed proportion of test statistics more (resp. at least as) extreme as the observed one. This strategy is further referred to as *full-empiric* and requires a large number of permutations to achieve a good precision in the estimation of small p-values. Here, we used 5000 permutations.

As an alternative, we propose a *semi-empiric* approach in which a limited number of resampled statistics are used to estimate parameters of the test statistic distribution. In the *semi-empiric* approach, a limited ($N = 500$) number of resampled statistics are used to estimate the mean (μ) and standard deviation ($\sigma$) of the test statistic under $H_0$. Assuming that the test statistic follows a normal distribution, we can then compute the p-value corresponding to an observed statistic by using the $\mathcal{N}(\mu, \sigma^2)$ distribution. For the likelihood ratio test (LRT) statistic, which asymptotically follows a chi-square distribution under the null, we transformed the statistic into a Z-score by taking the square root and signing according to the direction of effect. This approach allows a better precision in the p-value estimation, or permutation sparing, provided that the underlying hypothesis on the distribution is true.

A R script for the *LocPerm* procedure with a test example are available at https://www.iame-research.center/eq4/resources/ and at https://www.institutimagine.org/abel

#### c. *Rare variant association test*

A number of methods have been proposed to test for the association between rare variants and a phenotype which are based on the aggregation of rare variants within a genetic unit, e.g. protein coding genes. Among them, we used in this study the "cohort allelic sum test" (CAST) approach (2) which assumes that the presence of one rare variant is sufficient to cause the phenotype and is particularly well suited to study limited numbers of rare monogenic disorders which was the focus of our study. We implemented the CAST approach in the logistic regression framework and used the LRT statistic. As a reference method to account for population stratification we included the first three PCs of the PC analysis on common variants in the association model (denoted as CAST-3PC). We compared results obtained using 1) the asymptotic distribution of CAST and CAST-3PC, 2) standard permutations applied to CAST, and 3) *LocPerm* applied to CAST.

#### d. *Simulation study*

For the simulation study, we used two real NGS datasets (public and in-house), in order to have realistic site frequency spectrum and LD structure. The in-house dataset, referred as HGID (Human Genetic of Infectious Diseases Database), is composed of 3,104 WES data generated with the exome capture kit SureSelect Human All Exon V4+UTRs (https://agilent.com). The public dataset is composed of 2,504 whole-genomes from the 1000 genome phase 3 (http://www.internationalgenome.org/). We first merged the two datasets and extracted only the exonic regions captured by the Agilent V4+UTRs capture kit. We performed genotype and variant level quality control to focus only on high quality coding variants defined as having a depth of coverage (DP) > 8, a genotype quality (GQ) > 20, a minor read ratio (MRR) > 0.2 and a call-rate > 95% (3). We then excluded all related individuals based on the kinship coefficient (King's kinship 2K > 0.1875 (4,5)) leading to a total of 4,887 unrelated samples. From the whole dataset, we selected samples of European ancestry including all individuals with a reported European ancestry from 1000 genomes (i.e. CEU, TSI, FIN, GBR and IBS) and HGID cohorts. In addition, we included samples with unknown recorded ancestry from HGID but with a genetic distance to the randomly selected sample HG00146 form the 1000 genomes GBR population lower than the maximum genetic distance observed between HG00146 and other samples with known European ancestry. This led to final sample of 1523 European individuals.

We empirically separated this European sample in three parts according to their ancestry based on PC analysis (efigure 1): 127 individuals of Northern ancestry (mainly including the 1000 genomes FIN samples), 651 of Middle-Europe ancestry (including the 1000 genomes CEU and GBR samples) and 745 of Southern ancestry (including the 1000 genomes TSI and IBS samples). Cases and controls samples were simulated under three stratification scenario: 1) no stratification scenario where cases were equally distributed across the three sub-populations; 2) intermediate stratification scenario where 5/6 of cases came from Southern Europe, 1/6 from Middle Europe and Northern Europe (4:1 ratio); and 3) extreme stratification where all cases came from Southern Europe. In all scenarios, controls were equally distributed across the three sub-populations.

Under the null hypothesis of no genetic association, samples of 30, 60 or 120 cases and 60, 120 or 180 controls were randomly drawn from the source population according to the three stratification scenario. We also generated samples of 381 cases and 1142 controls representing the whole source population. For each configuration we generated 15 replicates and performed the analysis of all protein coding genes with at least 10 carriers of a rare variant defined as having a MAF ≤ 0.05 in the analyzed cohort. According to the sample size and the stratification scenario the number of genes tested varied from 135,054 to 265,365 (Table 1). The type I error rate at nominal level α was computed as the number of p-values equal or lower than α divided by the total number of genes tested over the 15 replicates.

For the power study, we selected the gene *RORC*, which contained 15 rare variants with a cumulative MAF of 6.2%, as being the disease-causing gene. Within *RORC*, we randomly selected 10 variants, with a cumulative MAF of 3%, as being disease-causing, leading to about 6% of the population carrying at least one risk allele. Individuals were considered as carrier if they had at least one risk allele and no cumulative effect was simulated. Penetrances for the carrier/non-carrier categories were calculated for each stratification scenarios based on the proportion of cases and controls, the frequency of carriers, and a relative risk of 4. We generated 500 replicates of samples of 30/120, 30/180 and 60/180 cases/controls under the three stratification scenarios. The power at nominal level α was computed as the number of p-values equal or lower than α divided by the total number of replicates (n=500).

### 2. Supplementary figure

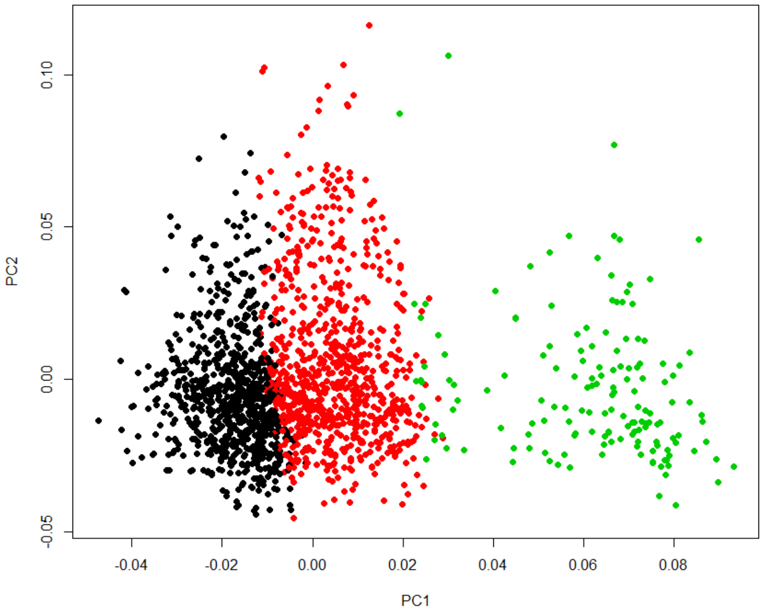

**eFigure 1:** Principal component representation of the source population (N=1523).

Black, red and green dots indicates individuals from Southern, Middle and Nothern Europe, respectively.

### 3. Supplementary table

**eTable 1: Type I error rate for different approaches and scenarios at a nominal α level of 5%.** Type I error rates above the upper bound of the 95% prediction interval in bold.

| Stratification | N cases | Ncontrols | N genes* | CAST | CAST-3PC | Std. Perm | LocPerm FE | LocPerm SE |
|---|---|---|---|---|---|---|---|---|
| Absence | 30 | 60 | 136932 | 5.3 | **5.9** | **5** | **5.1** | 4.9 |
| | 30 | 120 | 186513 | 6 | **6.4** | **4.9** | **5** | 5.3 |
| | 30 | 180 | 210287 | 5.9 | **6.5** | **4.9** | **4.9** | 5.1 |
| | 60 | 60 | 167621 | 5.6 | **5.7** | **5** | **5** | 5 |
| | 60 | 120 | 200883 | 5.3 | **5.5** | **4.9** | **5** | 4.9 |
| | 60 | 180 | 217518 | 5.3 | **5.5** | **4.9** | **5** | 5 |
| | 120 | 120 | 217319 | 5.4 | **5.5** | **5** | **5.1** | 5.1 |
| | 120 | 180 | 227650 | 5 | **5.3** | **4.9** | **4.9** | 4.9 |
| | 381 | 1142 | 265365 | 5.1 | **5.2** | **5** | **5** | 4.9 |
| Intermediate | 30 | 60 | 135896 | 6.3 | **6** | **5.9** | **5** | 4.8 |
| | 30 | 120 | 187282 | 7.2 | **6.2** | **5.9** | **5** | 4.8 |
| | 30 | 180 | 210510 | 6.9 | **6.3** | **5.8** | **4.9** | 4.7 |
| | 60 | 60 | 166912 | 7 | **5.4** | **6.2** | **5** | 4.7 |
| | 60 | 120 | 200578 | 7 | **5.6** | **6.6** | **5** | 4.6 |
| | 60 | 180 | 217720 | 7.2 | **5.6** | **6.8** | **4.9** | 4.6 |
| | 120 | 120 | 218007 | 7.9 | **5.3** | **7.4** | **5** | 4.7 |
| | 120 | 180 | 228317 | 8 | **5.2** | **7.8** | **4.8** | 4.4 |
| | 381 | 1142 | 265365 | 7.8 | **5.3** | **7.6** | **5.2** | 4.6 |
| Extreme | 30 | 60 | 135054 | 6.6 | **6.3** | **6.1** | **4.7** | 4 |
| | 30 | 120 | 187267 | 7.6 | **6.7** | **6.3** | **4.8** | 4.2 |
| | 30 | 180 | 210373 | 7.5 | **6.7** | **6.4** | **5** | 4.5 |
| | 60 | 60 | 167433 | 7.7 | **5.6** | **6.8** | **4.7** | 3.9 |

| | | | | | | | |
|---|---|---|---|---|---|---|---|
| 60 | 120 | 201217 | 7.6 | **5.8** | **7.2** | **4.8** | 4 |
| 60 | 180 | 218030 | 8 | **6** | **7.5** | **4.8** | 4.1 |
| 120 | 120 | 218063 | 8.9 | **5.3** | **8.4** | **4.8** | 4 |
| 120 | 180 | 228643 | 9.3 | **5.6** | **9** | **4.9** | 4.2 |
| 381 | 1142 | 265365 | 11.1 | **6** | **10.9** | **5** | 4.1 |

* N protein coding genes with at least 10 carriers of rare variants over 15 replicates
